# Submesoscale dynamics directly shape bacterioplankton community structure in space and time

**DOI:** 10.1101/2020.09.02.279232

**Authors:** Eduard Fadeev, Matthias Wietz, Wilken-Jon von Appen, Eva-Maria Nöthig, Anja Engel, Julia Grosse, Martin Graeve, Antje Boetius

**Affiliations:** Alfred Wegener Institute, Helmholtz Center for Polar and Marine Research, Bremerhaven, Germany; Max Planck Institute for Marine Microbiology, Bremen, Germany; GEOMAR Helmholtz Centre for Ocean Research, Kiel, Germany

**Keywords:** Arctic Ocean, phytoplankton bloom, submesoscale filaments, marginal ice zone, bacterioplankton, oceanic fronts

## Abstract

Submesoscale eddies and fronts are recognized as important components of oceanic mixing and energy fluxes. These submesoscale phenomena occur in the surface ocean for a period of a few days on scales between several hundred meters and a few tens of kilometers. Remote sensing and modeling suggest that they may influence marine ecosystem dynamics, but their limited temporal and spatial scales make them challenging for observation and in situ sampling. Here, the study of a submesoscale filament in summerly Arctic waters (depth 0 - 400 m) revealed enhanced vertical mixing of Polar and Atlantic water masses, resulting in a ca. 4 km wide and ca. 50 km long filament with distinct physical and biogeochemical conditions. Compared to the surrounding waters the filament was characterized by a distinct phytoplankton bloom dominated by diatoms and two-fold higher bacterioplankton cell densities. High-throughput 16S rRNA gene sequencing of both bacterioplankton communities revealed 3-4 orders of magnitude higher sequence abundance of *Synechococcus* inside the filament, as well as tenfold higher sequence abundance of taxonomic groups typically found during summertime in aging phytoplankton blooms (e.g., *Flavobacteriales*). In contrast, the surrounding waters contained severalfold higher sequence abundance of winterly taxonomic groups that are also associated with polar water masses (e.g., SAR202 clade). Altogether, our results show that physical submesoscale processes in the ocean can shape distinct biogeochemical conditions and microbial communities within a few kilometers. Furthermore, our results underline the importance of such submesoscale features for our understanding of surface ocean diversity and biogeochemical processes.

## Introduction

Microbial communities perform key functions in marine biogeochemical cycles by mediating dissolved and particular matter transformations and energy fluxes (Azam and Malfatti, 2007; Buchan et al., 2014). To date, regional biogeography of pelagic microbial communities is believed to be largely driven by large-scale (> 50 km) processes of physical mixing (Hewson et al., 2006). However, there is increasing evidence that mesoscale eddies and frontal systems have important ecological functions, featuring elevated phytoplankton growth (Lévy et al., 2001; Mouriño, 2004; Thomsen et al., 2016), unique bacterioplankton communities (Baltar et al., 2016; Djurhuus et al., 2017; Nelson et al., 2014; Zhang et al., 2011) and high microbial activity (Baltar et al., 2010; Baltar and Arístegui, 2017). Thus, physical oceanic processes on small spatial scales may have considerable effects on the composition and the activity of bacterioplankton communities, with probable influence on biogeochemical fluxes.

In recent years, observations by satellite remote sensing, autonomous profiling floats, and towed instruments revealed the ubiquity of submesoscale currents in the global ocean, confirming previous hypotheses based on numerical models (D’Asaro et al., 2011; Ferrari, 2011; Klymak et al., 2016; Lévy et al., 2012; Thomas et al., 2008; Thompson et al., 2016). These oceanic motions in surface waters are due to horizontal density gradients and take the shape of eddies and filaments with a lateral spatial range of a few hundred meters up to some tens of kilometers (Sasaki et al., 2014). Their motions persist for up to a few days and are characterized by strong lateral currents along the filament and an increased vertical circulation (McWilliams, 2016). The relatively small spatial scale and short-lived nature of such phenomena make them challenging for in situ biological observations (e.g., molecular sampling). To date, all existing observations from submesoscale filaments in the ocean have been focused on phyto- and zooplankton (reviewed in Lévy et al., 2018), and despite their relevance to processes in the ecosystem almost nothing is known regarding bacterioplankton communities in these filaments (Baltar et al., 2009, 2016; Baltar and Arístegui, 2017).

Investigating these dynamics in the Fram Strait is of importance considering its relevance for the ‘Atlantification’ of the Arctic Ocean and for other global climatic processes (Randelhoff et al., 2018; Wang et al., 2020). The Fram Strait connects the North Atlantic with the Arctic Ocean, and consists of two major surface current systems: (1) the East Greenland Current (EGC) flowing southwards along the Greenland shelf, transporting cold polar water and sea ice to the North Atlantic (de Steur et al., 2009); and (2) the West Spitsbergen Current (WSC) flowing northwards along the Svalbard Archipelago, transporting relatively warm and saline Atlantic water into the Arctic Ocean (Beszczynska-Möller et al., 2012; von Appen et al., 2016). The interactions between these distinct water masses result in a baroclinic instability in the central Fram Strait which facilitates the subduction of the Atlantic water below the polar water (Hattermann et al., 2016; von Appen et al., 2016; Wekerle et al., 2017). In addition to physicochemical differences between EGC and WSC, they have been shown to harbor distinct bacterioplankton communities (Fadeev et al., 2018; Müller et al., 2018; Wilson et al., 2017). Thus, the observed filament provides an excellent opportunity to investigate how bacterioplankton communities are shaped by submesoscale processes and how these patterns correspond to the general microbial distribution in the Fram Strait. Here we explored microbial dynamics within a submesoscale filament in the central Fram Strait (von Appen et al., 2018), the main gateway between the Atlantic and Arctic Oceans. Using high-throughput sequencing of the 16S rRNA gene we characterized free-living (0.22 - 3 μm) and particle-associated (> 3 μm) bacterial and archaeal communities (further addressed as bacterioplankton) inside and outside the filament, characterizing six depths from surface (10 m) to mesopelagic (400 m) waters. By combining molecular information with a set of biogeochemical parameters linked to a high-resolution physical analysis of this filament (von Appen et al., 2018), we identified differences in bacterioplankton community compositions between the filament and the surrounding waters, and investigated whether the observed differences are a result of mixing between water masses or internal succession within the communities. Specifically, we tested the hypothesis that bacterioplankton communities respond directly to the environmental conditions in submesoscale filaments and also shape them by their activities. These insights into the biological dynamics associated with such ubiquitous submesoscale features contribute to our understanding of their impact on ecosystem functioning in the global ocean.

## Results

### General characteristics of the submesoscale filament

In late July 2017 sea ice was present in the Fram Strait northeast and west of 0°EW/80°N (> 15 % ice concentration), while the eastern part of the Strait was under ice-free conditions (Figure 1A). On 26th of July 2017, satellite radar reflectivity image showed a nearly straight streak of sea ice in the marginal ice zone at 2.5°E/79°N, which extended over 50 km from northeast to southwest and had a width of only 500 m (Figure 1B). The high-resolution physical characterization of the phenomenon revealed two strong counteracting currents (change from +0.5m/s to −0.5m/s over 8km in NE-SW direction) in turbulent thermal wind balance with a distinct, a few kilometers wide filament of denser water below the observed sea-ice streak (von Appen et al., 2018). The surface ocean was strongly meltwater stratified. On either side of the filament, a meltwater layer was overlain by polar water (core depth ca. 30 m) which was in turn overlain by Atlantic water. By contrast, in the center of the filament meltwater was accumulated and overlain by Atlantic water which was located at a relatively shallow depth (Figure 1C). It was hypothesized that the denser Atlantic Water, which is typically present in the upper ocean in the West Spitsbergen Current further to the east, was in the process of subduction thereby releasing potential energy that drove the circulation (von Appen et al., 2018).

**Figure 1:**
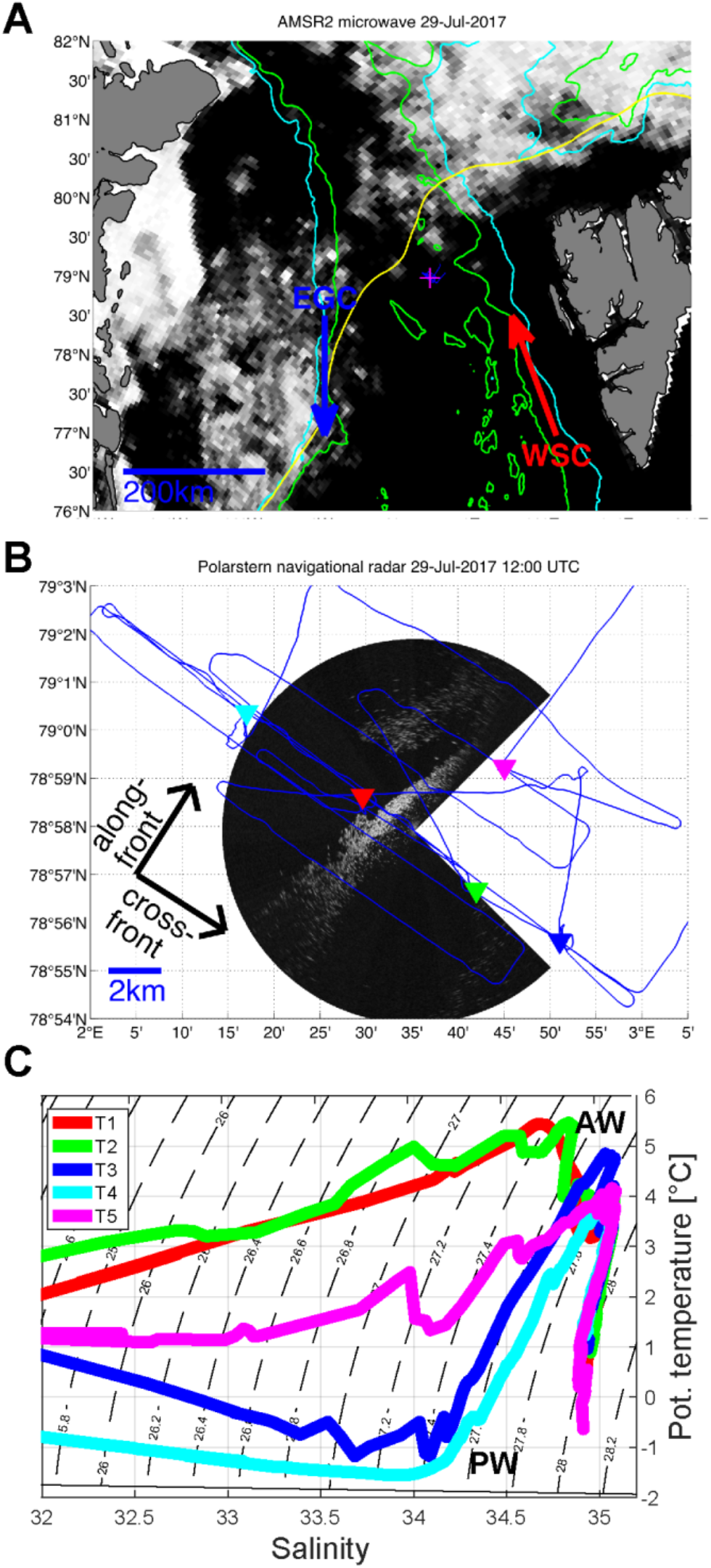
Sea ice distribution in the study area (black = open water, white = sea ice). (A) AMSR2 microwave sea ice concentration on 29 July 2017. Magenta plus marks the filament location. (B) RV Polarstern navigational radar reflectivity on 29 July 2017 12:00 UTC approaching the ice streak. Blue line shows the cruise track. Colored triangles mark the CTD stations: T1 (red), T2 (green), T3 (blue), T4 (cyan), and T5 (magenta). (C) TS diagram of the CTD profile at each sampling station, and the Atlantic Water (AW) and Polar Water (PW) characteristics. Adapted from von Appen et al. (2018).

### Biogeochemical conditions in the filament and the bounding currents

Based on previously defined temperature and salinity characteristics of the main water masses in the Fram Strait (Rudels et al., 2013), Atlantic waters temperature maximum > 5°C was observed inside the filament and Polar waters with temperature minima of ca. −1°C on both sides of it (Figure 1C). Water samples for biogeochemical and molecular analyses were collected at five stations: stations T1, T2 (profiles lacking polar water inside the filament), stations T3 and T4 (profiles with Polar Water in the bounding currents on either side), and station T5 (mixture of the two water masses at their intersection inside the filament; Figure 1). The horizontal distance between the stations was approximately 4 km. At each station, water samples were collected at six representative layers: surface (10 m), a chlorophyll *a* (chl-*a*) maximum (20 - 30 m), epipelagic (50 m and 100 m), and mesopelagic (200 m and 400 m).

At the time of sampling, nitrate (NO_3_) and phosphate (PO_4_) showed strong depletion in the upper 50 m of the water column inside the filament (stations T1, T2 and T5; Table 1). In surface waters (10 m) inside the filament the N:P ratios were 2.2-3.3, compared to much higher ratios (5.5-11.8) in the surface waters outside of the filament. In the upper 30 m, chl-*a* concentrations inside the filament were ranged between 0.6-2.9 mg m^−3^ with the highest values at T5 (Table 1). In contrast, outside of the filament chl-*a* concentrations were all below 1.0 mg m^−3^ with lowest values at T4. Concentrations of particulate organic carbon and biologically incorporated silica (bPSi) were also higher inside the filament (Table 1). Furthermore, partial pressure of carbon dioxide (pCO_2_) in surface waters inside the filament was below 250 μatm, compared to ca. 300 μatm outside of the filament.

**Table 1:**
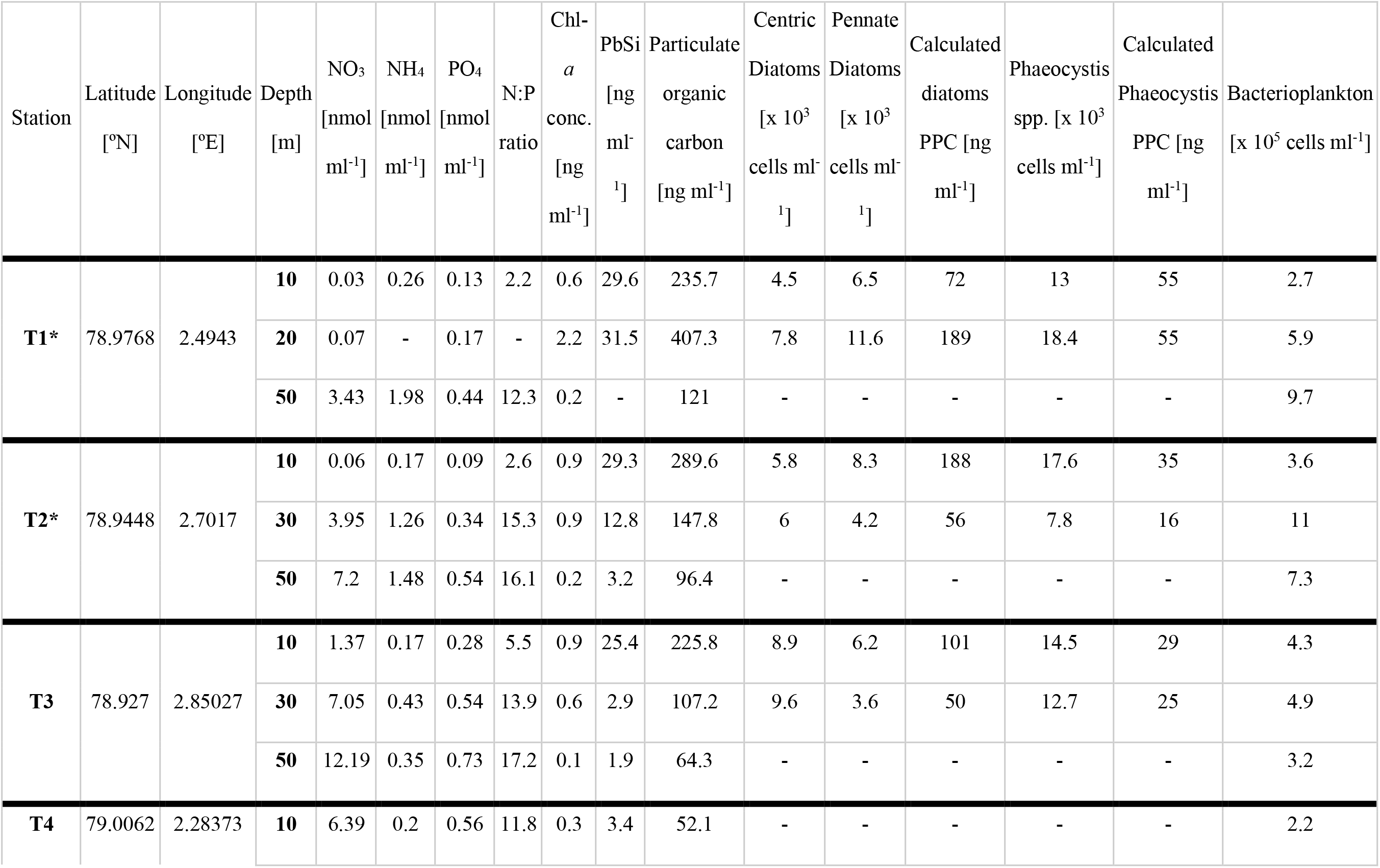
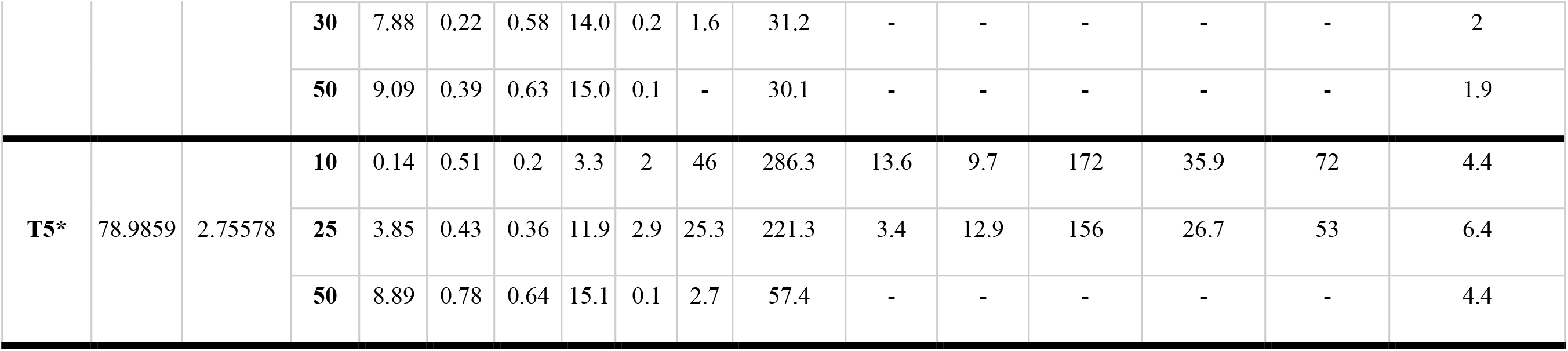
Overview of biogeochemical parameters and phytoplankton biomass in all sampled stations. The N:P ratio was calculated according to: (NO_3_+ NH_4_)/ PO_4_. (-) refers to no available data. (*) Stations inside the submesoscale filament.

In the upper 30 m, microscopic counts of major phytoplankton groups revealed higher cell densities of both *Phaeocystis* and diatoms inside the filament, with almost two-fold higher cell densities at T5 (Table 1). The estimated contribution of diatoms to the organic carbon budget (as a proxy for primary production) inside the filament was 50 % of the particulate organic carbon (Table 1).

### Abundance and diversity of bacterioplankton communities in the filament and the bounding currents

Cell density of the bacterioplankton communities correlated to the chl-*a* concentration pattern (Spearman’s rank correlation, r_s_ = 0.8, p < 0.001). Inside the filament bacterioplankton cell densities reached up to 1.1 x10^6^ cells ml^−1^ at T5. In contrast, bacterioplankton communities outside of the filament comprised two-fold lower densities, reaching values well below 0.5 x10^6^ cells ml^−1^ (Table 1). Below the upper 50 m depth, no differences in total cell densities were observed (Supplementary Table 1). At all stations and sampling depths, we characterized using 16S rRNA gene V4-V5 hypervariable region sequencing the free-living (FL) and the particle-associated (PA) bacterial and archaeal communities. The final dataset consisted of 5,770,041 sequences in 59 samples that were assigned to 3,980 amplicon sequence variants (ASVs; Supplementary Table 2). Rarefaction curves did not reach a plateau in any of the sampled communities, however, estimated asymptotic extrapolation to double amount of sequences showed only few additional ASVs (Supplementary Figure 2). Thus, our sequencing depth was satisfactory to represent most of the bacterioplankton diversity in all communities (Hsieh et al., 2018).

Alpha diversity estimators (observed ASVs, Chao1 richness and Shannon diversity index) showed statistically significant grouping of the bacterioplankton communities into two depth layers: upper 50 m (surface, chl-*a* max and upper epipelagic communities), and below 50 m (lower epipelagic and mesopelagic communities; ANOVA, p < 0.001; Tukey HSD, p < 0.01; Supplementary Figure 3). In the upper 50 m the number of observed ASVs and the Chao1 richness of the bacterioplankton communities were significantly lower inside the filament (t-test, p < 0.05). Below the upper 50 m, there were no significant differences between the alpha diversity of the bacterioplankton communities inside and outside the filament (t-test, p > 0.05; Figure 2A). Presence/absence analysis of the bacterioplankton communities (in both FL and PA fractions) revealed that in the upper 50 m 907 ASVs (23 % of total) were ubiquitous both inside and outside of the filament (Figure 2B). These ASVs were associated mainly with the bacterial order *Flavobacteriales* (161 ASVs) and the SAR11 clade (106 ASVs). Outside of the filament were 723 ASVs (18 % of total), associated mainly with the clades SAR202 (class *Dehalococcoidia*; 59 ASVs), SAR11 (56 ASVs; mainly ecotype II) and SAR406 (phylum *Marinimicrobia*; 53 ASVs), that were not present inside the filament. On the other hand, only 265 ASVs (7 % of total), comprised mainly of the bacterial order *Flavobacteriales* (43 ASVs) and the SAR11 clade (20 ASVs), were unique to the bacterioplankton communities inside the filament. Below the upper 50 m, 1885 ASVs (47 % of total) were observed in bacterioplankton communities both inside and outside of the filament, further 799 and 735 ASVs were unique to the bacterioplankton communities inside and outside the filament, respectively. All these ASVs were associated mainly with the order *Flavobacteriales* and the clades SAR11, SAR406 and SAR202.

**Figure 2:**
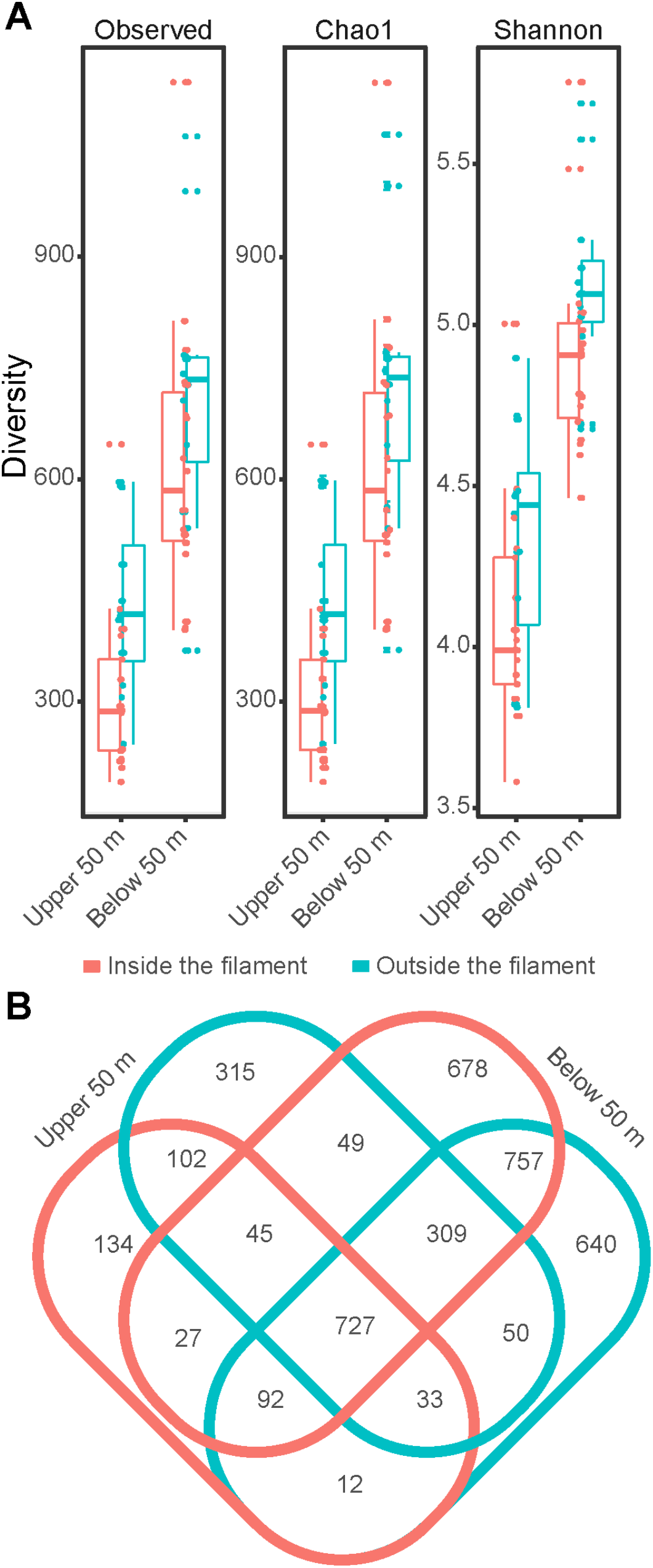
Diversity of bacterioplankton communities inside (red) and outside (blue) of the filament. (A) Alpha diversity indices of FL and PA communities divided between the upper 50 m and below them. (B) Venn diagram of shared and unique ASVs inside and outside of the filament.

### Taxonomic composition of bacterioplankton communities

In the upper 50 m, where most of the biogeochemical variability was observed, bacterioplankton communities also significantly differed in their composition inside and outside of the submesoscale filament (PERMANOVA, R^2^ = 0.13, p < 0.001; Figure 3A). Below the upper 50 m, there was no significant difference between the composition of the bacterioplankton communities inside and outside of the filament (PERMANOVA, p > 0.05). In order to investigate which taxonomic lineages account for the observed community differences inside and outside of the filament we performed differential ASV abundance tests in both FL and PA communities (Figure 3B). In the upper 50 m, 25 ASVs associated with the bacterial order *Flavobacteriales* (mainly families *Flavobacteriaceae* and *Cryomorphaceae*) and 23 ASVs of the SAR11 clade (mainly ecotype I and II), were significantly enriched inside the filament, with more than 10 times higher sequence abundance than outside of the filament (Figure 3B). Furthermore, inside the filament a single ASV associated with the cyanobacteria *Synechococcus* showed more than 4 orders of magnitude higher sequence abundance than outside of the filament (Supplementary Table 3). Outside of the filament the FL communities were significantly enriched by various taxonomic groups, such as the SAR202 clade (5 ASVs) and the archaeal order *Nitrosopumilales* (6 ASVs), reaching more than 10 times higher sequence abundance. The PA communities exhibited much smaller differences inside and outside of the filament (Figure 3B). Inside the filament the PA communities were significantly enriched by 10 ASVs of the family *Flavobacteriaceae*, with ca. 5 times higher sequence abundance than outside of the filament. Outside of the filament, the family *Pirellulaceae* (class *Planctomycetes*) was significantly enriched in the PA communities, with ca. 5 times higher sequence abundance than inside the filament.

**Figure 3:**
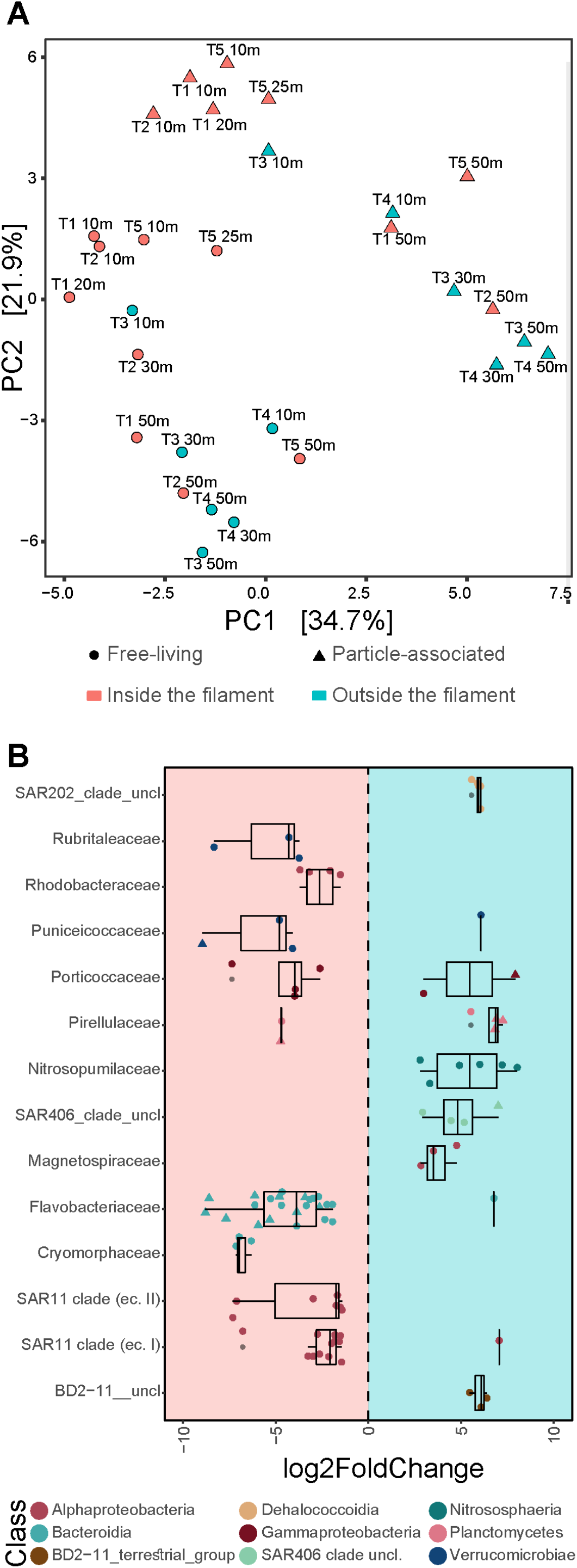
Overview of the bacterioplankton communities in the upper 50 m. (A) Principal component analysis (PCA) of free-living and particle-associated bacterioplankton communities based on Euclidean distances. The percentages on both axes represent the explained variance of the axis. (B) Differences in free-living (FL) and particle-associated (PA) community composition inside and outside of the submesoscale filament. The x axis represents the log_2_ fold change for all ASVs within a class. Positive value represents enrichment outside of the filament, and negative value represents enrichment in inside the filament. Different taxonomic classes are represented by color code. The shapes represent different fractions, circles free living communities and triangles particle-associated communities. Only taxonomic families with at least 3 significantly enriched (adjusted p value < 0.1) ASVs were included in the figure.

## Discussion

The biogeochemical and molecular comparison of surface seawater inside and outside of a submesoscale cyclonic filament that was observed in the Fram Strait (von Appen et al., 2018), revealed distinct biogeochemical conditions in the upper 50 m of the water column. In comparison to the neighboring waters, depleted inorganic nutrients and nutrient stoichiometry inside the filament, elevated concentrations of chl-*a*, drawdown of pCO_2_, and increase in particulate organic carbon may indicate that this was probably to a great extent due to phytoplankton growth. Microscopy counts of phytoplankton revealed slightly higher cell densities of both diatoms and *Phaeocystis* inside the filament in comparison to the surroundings suggesting an ongoing phytoplankton bloom inside the filament (Fadeev et al., 2018; Nöthig et al., 2015; Wu et al., 2007). Observations of blooms in low-latitude persistent submesoscale fronts confirm that these systems trigger growth a large fast growing phytoplankton, such as diatoms (Basterretxea and Arístegui, 2000; Allen et al., 2005; Li et al., 2012; Taylor et al., 2012; Clayton et al., 2014). Approx. 50% of the diatom population inside the submesoscale filament consisted of pennate diatoms. Pennate diatoms have been previously shown to bloom within only few days in iron fertilization experiments in the Southern Ocean (Coale et al., 1996; Landry et al., 2000). The increased numbers of *Phaeocystis* inside the filament compared to the surrounding waters may reflect their success in phytoplankton blooms in the Fram Strait in recent years (Engel et al., 2017; Nöthig et al., 2015). Taken together our results confirm previously established hypotheses that phytoplankton of high-latitudes can exploit submesoscale mixing processes and produce significant biomass within days, especially when warm waters are upwelled by fronts such as the observed one (Boyd et al., 2000; Taylor and Ferrari, 2011; Lévy et al., 2012, 2018).

Bacterioplankton showed some potential association with the different water masses, such as *Rhodobacterales* (class *Alphaproteobacteria*) with AW, and SAR202 (class *Dehalococcoidia*) with PW (Fadeev et al., 2018). However, the differences in the bacterioplankton communities inside and outside of the filament were mostly explained by the ongoing bloom inside the filament, especially at T5. In the upper 50 m we observed higher bacterioplankton cell densities and lower diversity of the communities inside the filament, suggesting selective growth of taxa. The bacterioplankton communities inside the filament were enriched (in terms of sequence abundance) by ASVs of the order *Flavobacteriales* (class *Bacteroidia*), that specializes on degrading complex organic biopolymers (Teeling et al., 2012; Buchan et al., 2014) and is typically found during phytoplankton blooms in the region (Wilson et al., 2017; Fadeev et al., 2018). Furthermore, we observed a strong enrichment of *Synechococcus* ASVs that suggests higher abundance of cyanobacteria inside the filament (Paulsen et al., 2016). On the other hand, outside of the filament the bacterioplankton communities had lower cell densities combined with higher diversity. These communities consisted of a large number of ASVs associated with the clades SAR406 (phylum *Marinimicrobia*) and SAR202 (class *Dehalococcoidia*), and were enriched by taxonomic groups such as the archaeal order *Nitrosopumilales*. All these taxonomic groups are typically found in the region’s surface polar waters and are representative of stable year-round or winter types, i.e. taxa not blooming in summer (Fadeev et al., 2018; Müller et al., 2018; Wilson et al., 2017).

The response of bacterioplankton communities to a phytoplankton bloom is often associated with elevated cell densities and a change in taxonomic composition (Bunse and Pinhassi, 2017). This shift reflects an enrichment of specialized taxonomic groups that utilize the phytoplankton-derived compounds (e.g., polysaccharides; Teeling et al., 2016). The high sequence proportion of ‘master recyclers’ (e.g., *Flavobacteriales*) together with high ammonia concentrations in the water suggest strong heterotrophic microbial activity inside the submesoscale filament. Previous comparison between various oceanic regions have shown that in water temperature higher than ca. 4°C there is no substantial difference in bacterial growth rates and activity between polar and low-latitude waters (Kirchman et al., 2009). Furthermore, previous measurements of bacterial activity in the Fram Strait (Piontek et al., 2014) showed comparable values to those measured in low-latitude frontal systems, and thus suggest potential turnover of the bacterioplankton communities on a scale of days at temperatures of 18-20 °C (Baltar et al., 2007). Taking into account the relatively warm temperatures in the upper 50 m inside the filament (up to 6°C in 30 m at station T2), we conclude that the bacterioplankton community inside the submesoscale filament went through an internal succession that followed the rapid development of the phytoplankton bloom. Thus, our results strongly indicate that bacterioplankton communities inside the submesoscale filament were shaped mainly by non-advective non-physical processes.

In conclusion, our observations confirm the hypothesis that submesoscale features can directly shape microbial diversity and community structure in the water column on a scale of few km and within a few days (Lévy et al., 2012, 2018). Despite their short temporal and spatial existence, the upwelling induced by submesoscale features promote phytoplankton blooms by nutrient supply from depth (Lévy et al., 2001; Mahadevan, 2015; Mouriño, 2004). The surplus of phytoplankton biomass then leads to a pronounced change in the abundance and composition of bacterioplankton communities inside the filament, towards dominance of ‘master recyclers’ and elevated microbial activity. However, despite the general similarity in the observed patterns, the submesoscale filament described here is different from previously studied frontal systems. In contrast to the frontal systems, that result from temperature gradients between constantly present two water masses (e.g., Baltar et al., 2009; Baltar et al., 2016), this submesoscale filament is a result of a strong salinity gradient between Polar and Atlantic waters close to the ice edge. The mixing between these water masses led to a formation of an enclosed water filament with distinct biogeochemical characteristics that persisted for several days (von Appen et al., 2018). Such strong salinity gradients can be found not only in the Fram Strait, but also in other sea-ice or freshwater-influenced regions worldwide (e.g., estuarine systems; Jaeger and Mahadevan, 2018). We suggest that the pronounced localized physical-biological phenomena that were described here play an important role in ocean productivity and export, in biogeochemical fluxes and biodiversity patterns of surface waters in large parts of the global ocean that features regular submesoscale processes.

## Material and Methods

### Water sampling and metadata collection

The sampling was carried out with 12 L Niskin bottles mounted on a CTD rosette (Sea-Bird Electronics Inc. SBE 911 plus probe) equipped with double temperature and conductivity sensors, a pressure sensor, altimeter, chlorophyll a fluorometer and transmissometer. Hydrographic data, including temperature and salinity are available at PANGAEA (von Appen and Rohardt, 2018). At all stations water samples were collected at five depths: 10m, 20 – 30m, 50m, 100m, 200m and 400m (Supplementary Table 2).

### Concentrations of inorganic nutrients

Inorganic nutrients were analyzed with a continuous flow autoanalyzer (Evolution III, Alliance Instruments, France). The unfiltered CTD samples were measured simultaneously on five channels: phosphate, silicate, nitrate, nitrite, and ammonium, essentially after Grashoff et al., 1983. The detection and quantification of ammonium was carried out using fluorescence spectroscopy after Kérouel and Aminot (1997). All measurements were calibrated with a five nutrients standard cocktail (all from Merck, traceable to SRM from NIST) diluted in artificial seawater (ASW), which was also used as wash-water between the samples. Each 20th run we checked our standards with Reference Material for Nutrients in Seawater (CRM 7602-a) produced by NMIJ. Standards and methods have been proven by inter calibration exercises (e.g., ICES and Quasimeme). Detection limit: c. 0.01 μmol l^−1^, Seawater Analysis after Grasshoff et al., 1983 (Verlag Chemie GmbH Weinheim). Detection limits: Nitrite, [NO_2_^−^], c. 0.002 μmol l^−1^; phosphate [PO_4_^3−^]_3_, c. 0.01 μmol l^−1^; silicate [Si(OH^−^)_4_], c. 0.03 μmol l^−1^; ammonium [NH4^+^], c. 0.01 - 0.02 μmol l^−1^.

The underway measurements of pCO_2_ were retrieved from the SOCAT database (socat.info), under expedition code 06AQ20170723.

### Concentrations of chlorophyll *a*, particulate organic carbon (POC) and biogenic silica (PbSi)

Concentrations of chlorophyll *a* (chl-*a*) were determined from 0.5 - 2 L of seawater filtered onto glass fiber filters (Whatman GF/F) under low vacuum pressure (< 200 mbar). Filters were stored at −20°C until analysis. Pigments were extracted with 10 ml of 90 % acetone, the filters treated with an ultrasonic device in an ice bath for less than a minute, and then further extracted in the refrigerator for 2 h. Subsequently filters were centrifuged for 10 minutes at 5000 rpm at 4°C prior to measurement. The concentration was determined fluorometrically (Turner Designs), together with total phaeophytin concentration after acidification (HCl, 0.1 N) based on methods described in Edler, 1979; Evans, 1980, respectively. The standard deviation of replicate test samples was < 10 %.

Particulate organic carbon (POC) was determined from 0.75 - 2 L of seawater filtered onto pre-combusted (4h at 500°C) glass fiber filters (Whatman GF/F) at low vacuum (<200 mbar). After filtration, the filters were stored frozen (−20°C) until analysis. Prior to analysis, filters were soaked in 0.1N HCl for removal of inorganic carbon and dried at 60°C. POC concentrations were determined with a CHN Elemental Analyzer by Carlo Erba (ThermoFisher Scientific, Rockford, IL, USA).

Particulate biogenic silica (PbSi) was determined following von Bodungen et al. (1991). Subsamples of 0.5 to 1 L were filtered on cellulose acetate filters (0.8 μm pore size), processed using the wet-alkaline method (with 0.2 M NaOH pretreated 12 h at 85°C in an oven), and extracted for 2 h at 85°C in a shaking water bath.

### Microscopic counts of phytoplankton groups

Seawater samples from 10 m and 20-30 m depths were preserved in hexamethylenetetramine-buffered formalin (final concentration 0.5 %) and stored in brown glass bottles. Phytoplankton cells were counted by light microscopy, respectively. For microscopic analyses an aliquot of 50 mL was transferred to settling chambers where cells were allowed to settle for 48 h. Phytoplankton cells were identified to genus level and at least 500 cells of the dominant phytoplankton species/groups were counted with an inverted microscope at three different magnifications using phase contrast.

Phytoplankton carbon content (PPC) was calculated by multiplying cell counts with the carbon value for the individual cells as determined following Edler (1979).

### Bacterioplankton cell densities

Cell densities were determined by flow cytometry (FACSCalibur, Becton Dickinson). Samples were fixed with glutaraldehyde at 1% final concentration and stored at −20°C. Prior to analysis, samples were stained with SybrGreen I (Invitrogen) before enumeration with visual inspection and manual gating of the populations in side scatter vs. green fluorescence cytograms. Fluorescent latex beads (Polyscience, Becton Dickinson) were used to normalize the counted events to volume (Gasol et al., 2000).

### DNA isolation and 16S rRNA amplicon sequencing

For assessing archaeal and bacterial community composition 2 - 4 L of seawater were filtered with a peristaltic pump (Masterflex; Cole Parmer) through successive membrane filters of 3 μm (Whatman Nucleopore, 47 mm polycarbonate), and 0.22 μm (Millipore Sterivex™ filters) pore sizes. All samples were stored at −20°C until DNA isolation. Genomic DNA was isolated from filter membranes to analyze particle-associated (PA) and free-living (FL) communities. The isolation was conducted by a combined chemical and mechanical procedure using the PowerWater DNA Isolation Kit (MO BIO Laboratories, Inc., Carlsbad, CA, USA). Prior to DNA isolation Sterivex™ cartridges were cracked open and filters transferred to kit-supplied bead beating tubes. The isolation was continued according to the manufacturer’s instructions, and DNA was stored at −20°C. Library preparation was performed according to the standard instructions of the 16S Metagenomic Sequencing Library Preparation protocol (Illumina, Inc., San Diego, CA, USA). The hyper variable V4–V5 region of 16S rRNA gene was amplified using primers 515F-Y (5’-GTGYCAGCMGCCGCGGTAA-3’) and 926R (5’-CCGYCAATTYMTTTRAGTTT-3’; Parada et al., 2016). Sequences were obtained on the Illumina HiSeq platform in a 2 × 250 bp paired-end run (CeBiTec Bielefeld, Germany) following the standard instructions of the 16S Metagenomic Sequencing Library Preparation protocol (Illumina, Inc., San Diego, CA, USA).

### Bioinformatics and statistical analyses

Raw paired-end reads were primer-trimmed using cutadapt (Martin, 2011). Further analyses were conducted using R v3.6.3 (http://www.R-project.org/) in RStudio v1.2.5033 (www.rstudio.org). Trimmed libraries were processed using DADA2 v1.14.1 (Callahan et al., 2016) following the suggested tutorial (https://benjjneb.github.io/dada2/tutorial.html). Briefly, chimeras and singletons were filtered and produced amplicon sequence variants (ASVs) taxonomically classified against Silva reference database release 138 (Quast et al., 2013). Taxonomically unclassified ASVs, ASVs assigned to bacterial or archaeal lineages, or assigned to mitochondria and chloroplast were excluded from further analysis.

Sample data matrices were managed using R package ‘phyloseq’ v1.28.0 (McMurdie and Holmes, 2013) and plots were generated using R package ‘ggplot2’ v3.3.0 (Gómez-Rubio, 2017). Rarefaction analysis was conducted using R package ‘iNEXT’ v2.0.20 (Hsieh et al., 2016). Calculation and visualization of shared ASVs was conducted using R package ‘venn’ v1.9 (Dussa, 2020). Statistical test were conducted using R package ‘vegan’ v2.5.6 (Oksanen et al., 2019). Prior to downstream analysis, a prevalence threshold (i.e., in how many samples did an ASV appear at least once) of 5% was applied on the ASV abundance table (McMurdie and Holmes, 2014). The fold-change in abundance of each ASV was calculated using the R package ‘DEseq2’ (v1.16.1; Love et al., 2014). The method applies a generalized exact binomial test on variance stabilized ASV abundance. The results were filtered by significance, after correction for multiple-testing according to Benjamini and Hochberg (1995) with an adjusted p-value < 0.1.

### Data Accession Numbers and Analyses Repository

Raw paired-end sequence, primer-trimmed reads were deposited in the European Nucleotide Archive under project accession number: PRJEB34666 (ENA; Silvester et al., 2018). The data were archived using the brokerage service of the German Federation for Biological Data (GFBio; Diepenbroek et al., 2014). Scripts for processing data can be accessed at ‘https://github.com/edfadeev/submesoscale_analysis’.

## Acknowledgments

We thank the captain and crew of RV Polarstern expedition PS107, as well as the chief scientist Ingo Schewe. Thanks to Christiane Lorenzen, Sandra Murawski and Nadine Knüppel for biochemical measurements, Malte Schmidt for microscopy counts of phyto- and protozooplankton communities, Tania Klüver for bacterioplankton counts using Flow Cytometry, and Halina Tegetmeyer for library preparation and sequencing of bacterioplankton samples. The authors also thank Federico Baltar for a critical review of the manuscript.

This work was conducted in the framework of the HGF Infrastructure Program FRAM of the Alfred-Wegener-Institute Helmholtz Center for Polar and Marine Research.

## Funding

This project has received funding from the European Research Council (ERC) under the European Union’s Seventh Framework Programme (FP7/2007-2013) research project ABYSS (grant agreement no. 294757) to AB. Microscopic counting was made available through CAO NERC/BMBF grant 03F0802B. Additional funding came from the Helmholtz Association, specifically for the FRAM infrastructure, from the Max Planck Society, and the Austrian Science Fund (FWF): M 2797-B.. Ship time was provided under grant AWI_PS107_07.

## Conflict of Interest Statement

The authors declare that the research was conducted in the absence of any commercial or financial relationships that could be construed as a potential conflict of interest.

## Notes

### Competing Interest Statement

The authors have declared no competing interest.

