## Supplemental Material for "Submesoscale dynamics directly shape bacterioplankton community structure in space and time"

### Submesoscale motions rapidly shape bacterioplankton communities in the Arctic Ocean

**Table 1:** Overview of phytoplankton and bacterioplankton cell densities in all sampled stations. (-) refers to no available data. * Stations inside the submesoscale filament.

| Sampling station | Depth [m] | Centric Diatoms [x $10^3$ cells $ml^{-1}$] | Pennate Diatoms [x $10^3$ cells $ml^{-1}$] | Phaeocystis spp. [x $10^3$ cells $ml^{-1}$] | Bacterioplankton [x $10^5$ cells $ml^{-1}$] |
|---|---|---|---|---|---|
| **T1*** | **10** | 4.5 | 6.5 | 13 | 2.7 |
| | **20** | 7.8 | 11.6 | 18.4 | 5.9 |
| | **50** | - | - | - | 9.7 |
| | **100** | - | - | - | 3.8 |
| | **200** | - | - | - | 2.3 |
| | **400** | - | - | - | 1.3 |
| **T2*** | **10** | 5.8 | 8.3 | 17.6 | 3.6 |
| | **30** | 6 | 4.2 | 7.8 | 11 |
| | **50** | - | - | - | 7.3 |
| | **100** | - | - | - | 4.4 |
| | **200** | - | - | - | 2.8 |
| | **400** | - | - | - | 1.9 |
| **T3** | **10** | 8.9 | 6.2 | 14.5 | 4.3 |
| | **30** | 9.6 | 3.6 | 12.7 | 4.9 |
| | **50** | - | - | - | 3.2 |
| | **100** | - | - | - | 2.5 |
| | **200** | - | - | - | 1.9 |
| | **400** | - | - | - | 0.6 |
| **T4** | **10** | - | - | - | 2.2 |
| | **30** | - | - | - | 2 |
| | **50** | - | - | - | 1.9 |
| | **100** | - | - | - | 3.3 |
| | **200** | - | - | - | 2.1 |
| | **400** | - | - | - | 1.1 |
| **T5*** | **10** | 13.6 | 9.7 | 35.9 | 4.4 |
| | **25** | 3.4 | 12.9 | 26.7 | 6.4 |
| | **50** | - | - | - | 4.4 |
| | **100** | - | - | - | 2.6 |
| | **200** | - | - | - | 2.0 |
| | **400** | - | - | - | 0.6 |

**Table 2**: Geographic origin and alpha diversity of free-living (FL) and particle-associated (PA) bacterioplankton communities at all sampled stations.

| PANGAEA sampling event | Latitude [º North] | Longitude [º East] | Station name | Depth [m] | Community | Total no. of sequences | No. of ASVs | Chao1 richness | Community completeness [%] | Shannon div. Index | Simpson div. Index | Pielou's evenness index |
|---|---|---|---|---|---|---|---|---|---|---|---|---|
| PS107/10-4 | 78.9768 | 2.4943 | T1 | 10 | FL | 97530 | 736 | 883.8 | 83 | 4.12 | 0.95 | 0.62 |
| PS107/10-4 | 78.9768 | 2.4943 | T1 | 10 | PA | 54831 | 586 | 752.1 | 78 | 3.68 | 0.94 | 0.58 |
| PS107/10-4 | 78.9768 | 2.4943 | T1 | 20 | PA | 112722 | 776 | 1001.6 | 77 | 4.05 | 0.96 | 0.61 |
| PS107/10-4 | 78.9768 | 2.4943 | T1 | 20 | FL | 94203 | 680 | 827 | 82 | 3.98 | 0.93 | 0.61 |
| PS107/10-4 | 78.9768 | 2.4943 | T1 | 50 | FL | 102786 | 871 | 1116.3 | 78 | 4.12 | 0.94 | 0.61 |
| PS107/10-4 | 78.9768 | 2.4943 | T1 | 50 | PA | 59237 | 653 | 892.6 | 73 | 3.99 | 0.96 | 0.62 |
| PS107/10-4 | 78.9768 | 2.4943 | T1 | 100 | FL | 98877 | 1741 | 2047.3 | 85 | 5.05 | 0.98 | 0.68 |
| PS107/10-4 | 78.9768 | 2.4943 | T1 | 100 | PA | 47813 | 1331 | 1779.5 | 75 | 4.85 | 0.98 | 0.67 |
| PS107/10-4 | 78.9768 | 2.4943 | T1 | 200 | FL | 110392 | 2234 | 2560.6 | 87 | 5.34 | 0.98 | 0.69 |
| PS107/10-4 | 78.9768 | 2.4943 | T1 | 200 | PA | 50299 | 1773 | 2203.1 | 80 | 5.23 | 0.98 | 0.7 |
| PS107/10-4 | 78.9768 | 2.4943 | T1 | 400 | FL | 51404 | 1923 | 2225.3 | 86 | 5.47 | 0.98 | 0.72 |
| PS107/10-4 | 78.9768 | 2.4943 | T1 | 400 | PA | 33237 | 1600 | 1982 | 81 | 5.41 | 0.98 | 0.73 |
| PS107/12-3 | 78.9448 | 2.7017 | T2 | 10 | PA | 67939 | 613 | 770 | 80 | 3.7 | 0.95 | 0.58 |
| PS107/12-3 | 78.9448 | 2.7017 | T2 | 10 | FL | 59012 | 611 | 739.2 | 83 | 4.05 | 0.95 | 0.63 |
| PS107/12-3 | 78.9448 | 2.7017 | T2 | 30 | FL | 78285 | 872 | 1148.1 | 76 | 4.31 | 0.95 | 0.64 |
| PS107/12-3 | 78.9448 | 2.7017 | T2 | 50 | PA | 61008 | 1131 | 1535.6 | 74 | 4.46 | 0.97 | 0.63 |
| PS107/12-3 | 78.9448 | 2.7017 | T2 | 50 | FL | 106579 | 1557 | 2112.8 | 74 | 4.71 | 0.97 | 0.64 |
| PS107/12-3 | 78.9448 | 2.7017 | T2 | 100 | PA | 33389 | 1386 | 1799.5 | 77 | 5.26 | 0.98 | 0.73 |
| PS107/12-3 | 78.9448 | 2.7017 | T2 | 100 | FL | 276537 | 2146 | 2353 | 91 | 5.05 | 0.98 | 0.66 |

| | | | | | | | | | | | | |
|---|---|---|---|---|---|---|---|---|---|---|---|---|
| PS107/12-3 | 78.9448 | 2.7017 | T2 | 200 | PA | 91483 | 2063 | 2532.7 | 81 | 5.23 | 0.98 | 0.69 |
| PS107/12-3 | 78.9448 | 2.7017 | T2 | 200 | FL | 224854 | 2234 | 2466.1 | 91 | 5.23 | 0.98 | 0.68 |
| PS107/12-3 | 78.9448 | 2.7017 | T2 | 400 | PA | 43884 | 1817 | 2322.1 | 78 | 5.35 | 0.98 | 0.71 |
| PS107/12-3 | 78.9448 | 2.7017 | T2 | 400 | FL | 114047 | 2024 | 2257.3 | 90 | 5.2 | 0.98 | 0.68 |
| PS107/14-1 | 78.927 | 2.85027 | T3 | 10 | PA | 141537 | 846 | 993.9 | 85 | 3.9 | 0.96 | 0.58 |
| PS107/14-1 | 78.927 | 2.85027 | T3 | 10 | FL | 75795 | 728 | 913 | 80 | 3.96 | 0.95 | 0.6 |
| PS107/14-1 | 78.927 | 2.85027 | T3 | 30 | PA | 109210 | 1293 | 1715.5 | 75 | 4.41 | 0.97 | 0.61 |
| PS107/14-1 | 78.927 | 2.85027 | T3 | 30 | FL | 134277 | 1491 | 1817.5 | 82 | 4.64 | 0.97 | 0.64 |
| PS107/14-1 | 78.927 | 2.85027 | T3 | 50 | PA | 28972 | 1232 | 1623.3 | 76 | 5.21 | 0.98 | 0.73 |
| PS107/14-1 | 78.927 | 2.85027 | T3 | 50 | FL | 125404 | 1889 | 2190.7 | 86 | 5.23 | 0.98 | 0.69 |
| PS107/14-1 | 78.927 | 2.85027 | T3 | 100 | PA | 93429 | 1980 | 2440.5 | 81 | 5.39 | 0.98 | 0.71 |
| PS107/14-1 | 78.927 | 2.85027 | T3 | 100 | FL | 160174 | 2169 | 2487.5 | 87 | 5.34 | 0.99 | 0.69 |
| PS107/14-1 | 78.927 | 2.85027 | T3 | 200 | PA | 83642 | 1896 | 2254.1 | 84 | 5.3 | 0.98 | 0.7 |
| PS107/14-1 | 78.927 | 2.85027 | T3 | 200 | FL | 277864 | 2318 | 2482.2 | 93 | 5.42 | 0.98 | 0.7 |
| PS107/14-1 | 78.927 | 2.85027 | T3 | 400 | PA | 27452 | 1657 | 1918.3 | 86 | 5.94 | 0.99 | 0.8 |
| PS107/16-3 | 79.0062 | 2.28373 | T4 | 10 | PA | 45769 | 871 | 1158.5 | 75 | 4.01 | 0.93 | 0.59 |
| PS107/16-3 | 79.0062 | 2.28373 | T4 | 10 | FL | 160729 | 1635 | 2014.4 | 81 | 4.61 | 0.96 | 0.62 |
| PS107/16-3 | 79.0062 | 2.28373 | T4 | 30 | PA | 53191 | 1202 | 1569.5 | 77 | 4.35 | 0.95 | 0.61 |
| PS107/16-3 | 79.0062 | 2.28373 | T4 | 30 | FL | 101950 | 1536 | 1846.6 | 83 | 4.72 | 0.97 | 0.64 |
| PS107/16-3 | 79.0062 | 2.28373 | T4 | 50 | PA | 46763 | 1399 | 1864.1 | 75 | 4.67 | 0.97 | 0.64 |
| PS107/16-3 | 79.0062 | 2.28373 | T4 | 50 | FL | 113568 | 1847 | 2133.9 | 87 | 4.96 | 0.97 | 0.66 |
| PS107/16-3 | 79.0062 | 2.28373 | T4 | 100 | PA | 85030 | 1491 | 1890.7 | 79 | 4.87 | 0.98 | 0.67 |
| PS107/16-3 | 79.0062 | 2.28373 | T4 | 100 | FL | 100314 | 1825 | 2086.7 | 87 | 5.2 | 0.98 | 0.69 |

| | | | | | | | | | | | | |
|---|---|---|---|---|---|---|---|---|---|---|---|---|
| PS107/16-3 | 79.0062 | 2.28373 | T4 | 200 | PA | 58967 | 1654 | 1903.7 | 87 | 5.4 | 0.98 | 0.73 |
| PS107/16-3 | 79.0062 | 2.28373 | T4 | 200 | FL | 152855 | 2147 | 2452.2 | 88 | 5.37 | 0.98 | 0.7 |
| PS107/16-3 | 79.0062 | 2.28373 | T4 | 400 | PA | 63864 | 1618 | 2021.1 | 80 | 5.39 | 0.98 | 0.73 |
| PS107/16-3 | 79.0062 | 2.28373 | T4 | 400 | FL | 103741 | 2604 | 2872.6 | 91 | 6 | 0.99 | 0.76 |
| PS107/18-3 | 78.9859 | 2.75578 | T5 | 10 | PA | 29673 | 508 | 706.2 | 72 | 3.91 | 0.95 | 0.63 |
| PS107/18-3 | 78.9859 | 2.75578 | T5 | 10 | FL | 181574 | 875 | 1092.3 | 80 | 4.11 | 0.95 | 0.61 |
| PS107/18-3 | 78.9859 | 2.75578 | T5 | 25 | PA | 72662 | 809 | 1122.2 | 72 | 4.17 | 0.97 | 0.62 |
| PS107/18-3 | 78.9859 | 2.75578 | T5 | 25 | FL | 191066 | 1129 | 1325.3 | 85 | 4.39 | 0.96 | 0.62 |
| PS107/18-3 | 78.9859 | 2.75578 | T5 | 50 | PA | 72894 | 1059 | 1390.3 | 76 | 4.54 | 0.97 | 0.65 |
| PS107/18-3 | 78.9859 | 2.75578 | T5 | 50 | FL | 140290 | 2067 | 2433.8 | 85 | 5.36 | 0.99 | 0.7 |
| PS107/18-3 | 78.9859 | 2.75578 | T5 | 100 | PA | 63608 | 1338 | 1659.5 | 81 | 5.12 | 0.98 | 0.71 |
| PS107/18-3 | 78.9859 | 2.75578 | T5 | 100 | FL | 91629 | 2296 | 2625.7 | 87 | 5.53 | 0.99 | 0.71 |
| PS107/18-3 | 78.9859 | 2.75578 | T5 | 200 | PA | 29345 | 1095 | 1418.9 | 77 | 5.09 | 0.98 | 0.73 |
| PS107/18-3 | 78.9859 | 2.75578 | T5 | 200 | FL | 62544 | 2219 | 2671.2 | 83 | 5.6 | 0.99 | 0.73 |
| PS107/18-3 | 78.9859 | 2.75578 | T5 | 400 | FL | 89674 | 2346 | 2659.2 | 88 | 5.98 | 0.99 | 0.77 |
| PS107/18-3 | 78.9859 | 2.75578 | T5 | 400 | PA | 43165 | 863 | 918.1 | 94 | 5.38 | 0.99 | 0.8 |

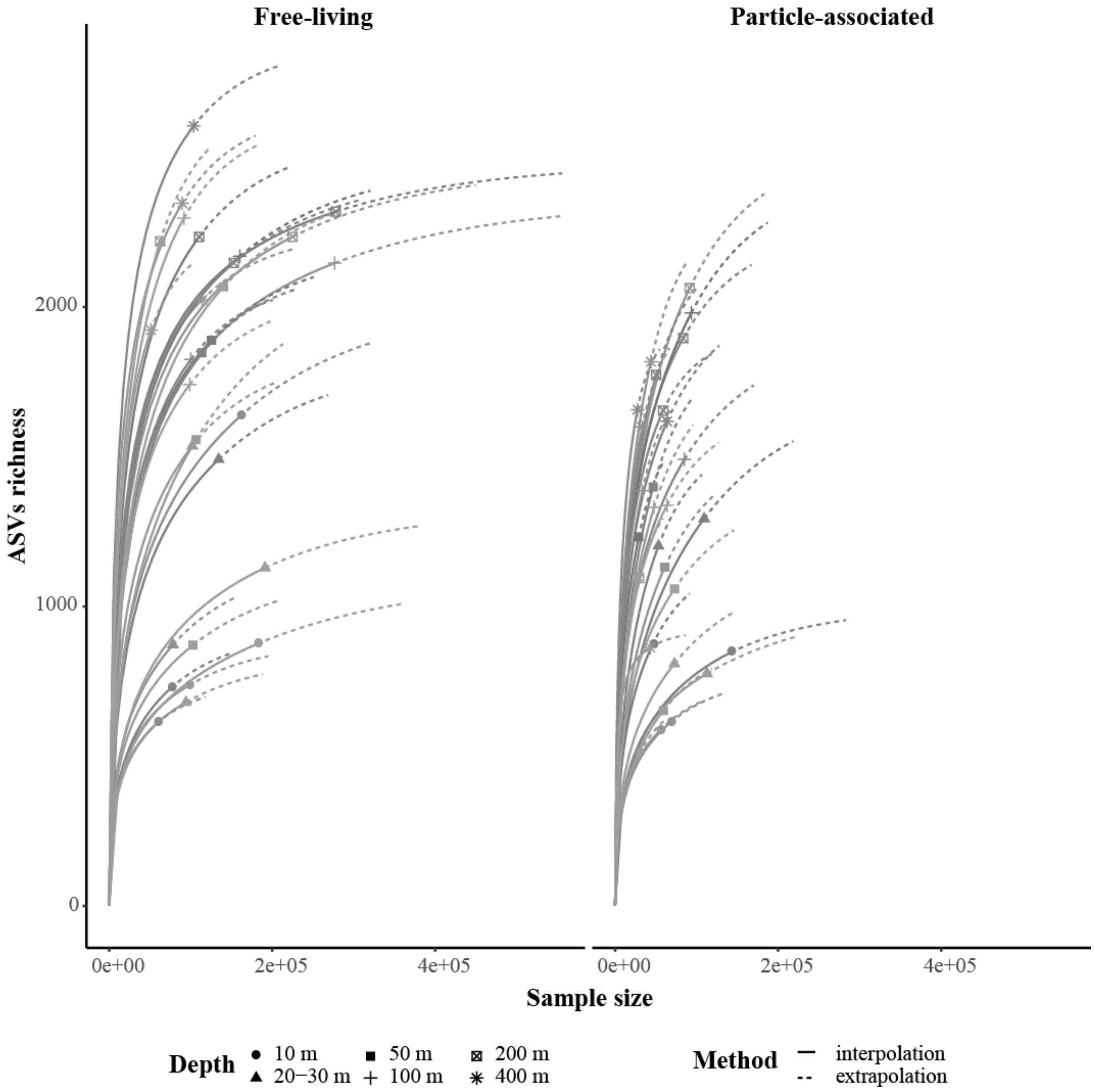


**Figure 1:** Rarefaction analysis of bacterioplankton 16S rRNA gene amplicon sequences. Solid lines represent the observed accumulation of ASVs (y axis) with the number of reads sampled (x axis), and dashed lines the extrapolated accumulation up to the double amount of reads. The observed values for each community are denoted by solid shapes. Sample-size-based rarefaction curves generated with the R-package "iNEXT", based on the Hill number of order q = 0. The rarefaction curves for each sample were generated based on 40 equally spaced rarefied sample sizes with 100 iterations.

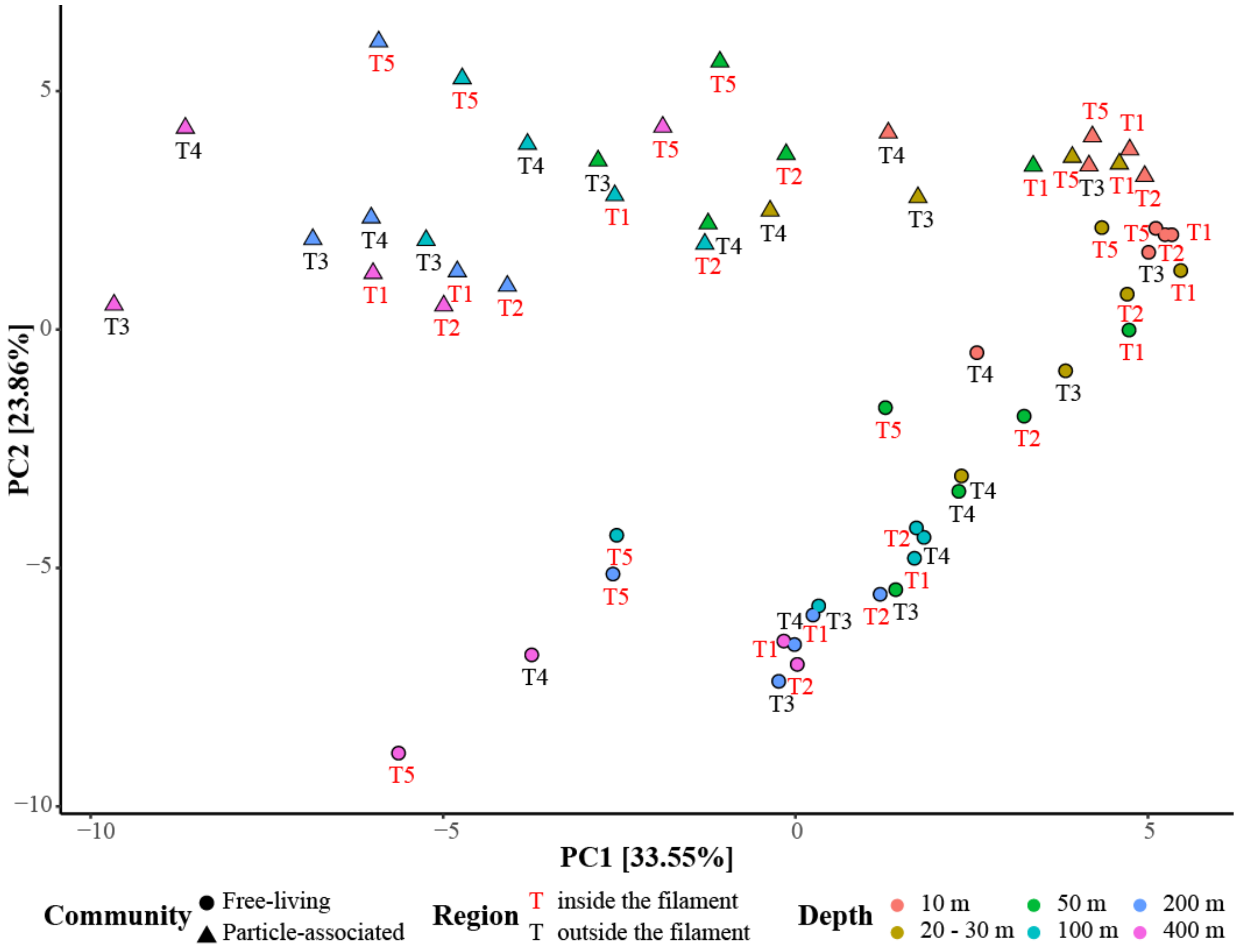


**Figure 2:** Principal component analysis (PCA) of free-living and particle-associated bacterioplankton communities based on Euclidean distances. The percentages on both axes represent the explained variance of the axis.

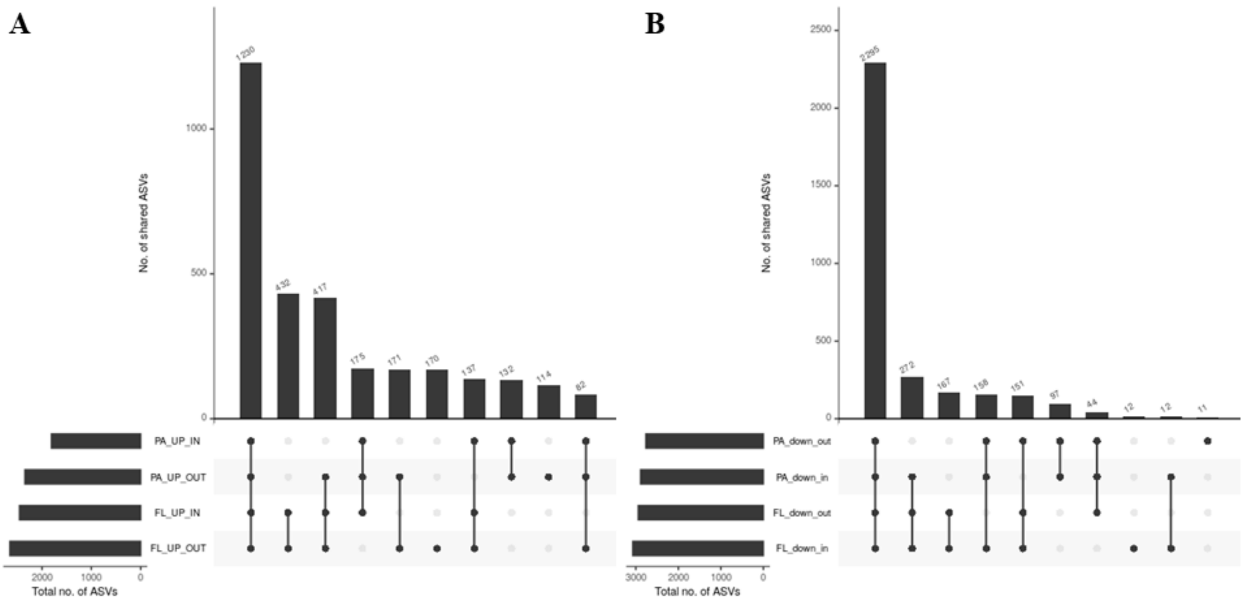


**Figure 3:** Comparison of community composition between the different regions and fractions, in the upper 50 m (**A**) and in 50-400 m depth (**B**). The main y-axis represent the number of shared ASVs between the different groups of communities. FL and PA represent the different fractions, in- inside the filament and out-outside of the filament.
